# Multi-antigen recognition circuits overcome challenges of specificity, heterogeneity, and durability in T cell therapy for glioblastoma

**DOI:** 10.1101/2021.01.07.425632

**Authors:** Joseph H. Choe, Payal B. Watchmaker, Milos S. Simic, Ryan D. Gilbert, Aileen W. Li, Nira A. Krasnow, Diego A. Carrera, Wei Yu, Kira M. Downey, Anna Celli, Juhyun Cho, Jessica D. Briones, Ruth Dannenfelser, Lia Cardarelli, Sachdev S. Sidhu, Kole T. Roybal, Hideho Okada, Wendell A. Lim

## Abstract

Treatment of solid cancers with chimeric antigen receptor (CAR) T cells is plagued by the lack of target antigens that are both tumor-specific and homogeneously expressed. We show that multiantigen prime-and-kill recognition circuits have the flexibility and precision to overcome these challenges in attacking glioblastoma. A synNotch receptor that recognizes a specific priming antigen – the heterogeneous glioblastoma neoantigen EGFRvIII or a brain tissue-specific antigen – is used to locally induce expression of a CAR, enabling thorough but controlled tumor killing by targeting of homogeneous antigens that are not absolutely tumor specific. Moreover, regulated CAR expression maintains a higher fraction of the T cells in the naïve-like state which is associated with higher durability *in vivo*. In summary, using circuits that integrate recognition of multiple imperfect but complementary antigens, we improve the specificity and persistence of T cells directed against glioblastoma, providing a general recognition strategy applicable to other solid tumors.

---

Although chimeric antigen receptor (CAR) T cells have demonstrated remarkable outcomes in treating hematologic malignancies (*1*), development of effective CAR T therapies for solid cancers remains a challenge, in large part due to the difficulty in identifying optimal target surface antigens. Very few antigens are truly tumor-specific, and on-target/off-tumor cross-reaction with normal tissues can cause lethal toxicities (*2-5*). Moreover, even if highly tumor-specific antigens are identified, these targets are often heterogeneously expressed, and selective CAR targeting allows for escape of antigen-negative tumor cells (*6*). Thus, there is a general need for novel tumor recognition strategies that can navigate the dual challenges of specificity and heterogeneity to enlarge the therapeutic window for safely and effectively attacking solid cancers.

A concrete example of this dual challenge is found in glioblastoma (GBM) (**Fig. 1A**). The epidermal growth factor (EGFRvIII) is a highly GBM-specific neoantigen found in a subset of GBM patients (*7-10*). But in prior clinical studies, targeting GBM with an EGFRvIII CAR led to tumor recurrence – the high heterogeneity of EGFRvIII expression allowed escape of EGFRvIII^-^ tumor cells, despite effective killing of EGFRvIII^+^ cells (*6, 11, 12*). In contrast, alternative glioma-associated surface antigens, including Ephrin type A receptor 2 (EphA2) and IL13 receptor α2 (IL13Rα2), are expressed in the vast majority of GBM cells (*13-15*), but have imperfect specificity. While they are not expressed in normal brain tissues, they are expressed at low levels in some non-tumor, non-brain tissues (*13-15*). In summary, it is challenging to find a single ideal surface GBM-antigen that is both specific and homogeneous.

**Figure 1.**
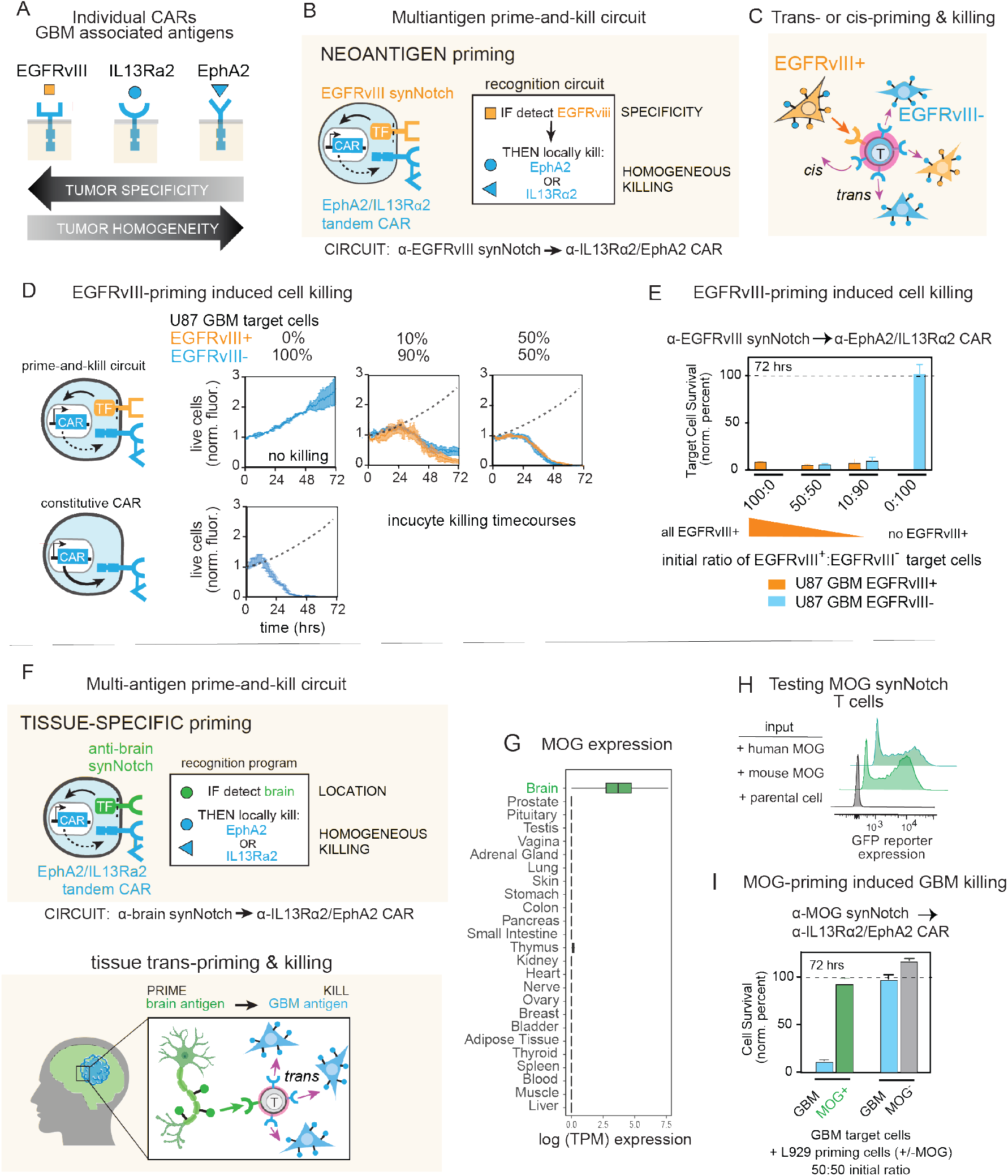
Multi-antigen prime-and-kill T cell circuits provide general strategy to maintain high tumor recognition specificity while overcoming antigen heterogeneity. A. Limitations of targeting GBM using single-antigen CAR T cells. Neoantigen EGFRvIII is highly tumor-specific, but heterogeneous, allowing tumor escape from anti-EGFRvIII CAR T cell treatment (*6, 11, 12*). Conversely, other potential target antigens, like EphA2 or IL13Rα2, are more homogeneously expressed in GBM, but are also expressed in some normal tissues, leading to potential on-target, off-tumor toxicity. The dual challenges of heterogeneity and specificity inherently limit the therapeutic window for CAR T cells. B. Design of synNotch→CAR circuit priming on EGFRvIII neoantigen. α-EGFRvIII synNotch receptor induces expression of tandem α-IL13Rα2/EphA2 CAR (4-1BBζ) (see **Fig. S1a**). This circuit harnesses the specificity of EGFRvIII and the homogeneous expression of the tandem killing antigens. C. The prime-and-kill CAR T cells should only be activated to kill EphA2+ or IL13Rα2+ target cells when exposed to EGFRvIII+ cells. These T cells should in principle be capable of cis- or trans-priming/killing. In cis-priming/killing, the priming and killing antigens are expressed on the same cell, while in trans-priming/killing these antigens are expressed on different cells. D. Time-course of killing assays using different heterogeneous mixtures of EGFRvIII+/EGFRvIII-target cells show that efficient trans-priming/killing can take place. Primary CD8+ human T cells with the α-EGFRvIII synNotch→ α-IL13Rα2/EphA2 CAR (or α-IL13Rα2/EphA2 constitutive CAR) were cultured with the indicated target cell mixtures (U87 cells engineered with or without EGFRvIII) at an E:T ratio of 5:1 and imaged over 3 days using an IncuCyte system. Tracking of EGFRvIII+ cell population (priming cells) is shown in yellow, while tracking of EGFRvIII-cell population (target cells) is shown in blue. Dotted black line shows the growth of 100% target cells as a reference (n=3, error bars are SEM). E. Summary of relative cell-type survival in experiments from panel D (at 72 hours). Killing of all target cells can be efficiently primed with a ratio of priming cells (EGFRvIII+) of as low as 10%; no killing, however, is observed in absence of priming cells (n=3, error bars are SD). F. Design of synNotch→CAR circuit that is primed by brain-specific antigen (“tissue-specific priming” circuit). Here a brain-specific synNotch receptor induces expression of tandem α-IL13Rα2/EphA2 CAR (4-1BBζ) (see **Fig. S1a**). Such a circuit is designed to limit killing only to the brain, preventing attack of normal tissues (non-brain) that express the killing antigens (e.g. EphA2 and IL13Rα2 are expressed in normal tissues outside of brain). G. MOG is a candidate brain-specific priming antigen. Box and whisker plots showing tissue specific expression of MOG across a subset of tissue samples in GTEx v7. Units shown are log scaled normalized RNAseq counts (Transcripts Per Million) taken from GTEx portal v7 (https://gtexportal.org/). See **Fig S7A** for analysis of another candidate brain-specific antigen (CDH10). H. Engineering of α-MOG synNotch receptor. Primary CD8+ T cells expressing α-MOG synNotch and GFP reporter were co-cultured with either parental K562 or K562 transduced to express mouse or human MOG. FACS histograms show induction of GFP reporter only in the presence of mouse or human MOG+ K562 and not K562 parental cells (representative of 3 experiments). I. Killing experiments shows that efficient brain antigen induced trans-priming/killing can take place *in vitro*. Primary CD8+ T cells with α-MOG synNotch → α-IL13Rα2/EphA2 CAR circuit were co-cultured with GBM target cells (GBM6) and L929 priming cells either expressing or not expressing mouse MOG. Relative cell survival of both target and priming cells was tracked over 72 hours. Experiments showed ability of these T cells to kill GBM6 target cells, but only in the presence of MOG+ priming cells (n=3, error bars are SD). Significant cytotoxicity in the GBM6 co-cultured with L929 mouse MOG was observed (**** p < 0.0001; t test) compared to the GBM6 co-cultured with L929 Parental cells. Cell population ratio: 1:1:1, 1e4 cells each. No significant killing of the L929 priming cell population is observed.

T cells that recognize multi-antigen combinations provide a possible solution to problems of antigen heterogeneity and specificity. We previously developed “prime-and-kill” circuits in which a synNotch receptor (an engineered receptor that activates a transcriptional output when it recognizes its cognate antigen; (*16*), induces expression of a CAR directed against a killing antigen (*17, 18*). Here we hypothesized that, by carefully choosing the priming and killing antigens (the synNotch and CAR ligands, respectively), such circuits could lead to hybrid recognition behaviors that might provide a way to navigate the tradeoffs between antigen specificity and heterogeneity.

We pursued two different strategies – priming with either a tumor-specific but heterogeneous antigen, such as EGFRvIII (**Fig. 1B-E**) or priming with a brain-specific antigen, such as myelin oligodendrocyte glycoprotein (MOG) (**Fig. 1F-H**). Priming by these antigens was then used to locally induce expression of a CAR that recognizes the more homogeneous antigens, EphA2 and IL13Rα2. The imperfect tumor-specificity of EphA2 and IL13Rα2, makes them non-ideal targets for conventional, single-target CAR T cell therapy approaches. These antigens could, however, serve as effective killing targets, if higher tumor selectivity was provided by the priming antigen. We hypothesized that T cells engineered with prime-and-kill circuits could induce local CAR-driven cytotoxicity restricted only to the vicinity of the priming cells, thereby avoiding indiscriminate killing in distant normal tissues that express the killing antigen but that lack the priming antigen. This type of circuit spatially integrates recognition of two imperfect but complementary antigen targets: the priming antigen provides specificity, while the killing antigen ensures homogeneity of the therapeutic attack. In all of these circuits, to achieve homogeneous killing and to further reduce the possibility of tumor escape, we utilized a tandem CAR that simultaneous targets the two killing antigens, EphA2 OR IL13Rα2 (*13, 19*). The tandem CAR, which functions as an OR gate, has an extracellular region containing an α-EphA2 single chain antibody and a IL13 mutein (variant of IL13 ligand that binds with higher affinity to IL13α2 over IL13Rα1) (*19, 20*) (see **Fig. S1a** for details of tandem CAR design).

In order for this strategy to overcome heterogeneity, a key question is whether a prime-and-kill T cell primed by one cell (EGFRvIII^+^) can then kill a different, neighboring target cell (EGFRvIII^-^) – a process we term *trans-priming/killing* (**Fig. 1C**) (*cis-priming/killing* would describe priming and killing based on antigens presented on the same cell). To first test for trans-killing, we used U87 “priming” GBM cells that were engineered to stably express the priming antigen EGFRvIII, and native U87 “target” cells (which endogenously express both EphA2 and IL13Rα2 killing antigens but are negative for EGFRvIII) (*11, 12, 21, 22*). We mixed the U87-EGFRvIII^+^ and U87-EGFRvIII^-^ cells in varying ratios to recapitulate different levels of heterogeneity observed in GBM patients (10-100% priming cells) then tested if target cell killing was induced by the presence of priming cells (**Fig. 1D, E**).

We found that CD8+ T cells engineered with the **α-EGFRvIII synNotch** → **α-IL13Rα2/EphA2 CAR** prime-and-kill circuit can effectively kill EGFRvIII^-^ targets cells *in vitro*, even with as low as 10% EGFRvIII^+^ priming cells (**Fig. 1D, E**). In contrast, no killing of the EGFRvIII^-^ target cells was observed in the absence of priming cells. In these assays, we tracked the kinetics of killing of the two different tumor cell populations (EGFRvIII- and EGFRvIII+) over 72 hours (see **Movie S1** for time-lapse of killing). Effective killing was observed with as low as 10% priming cells, although killing was slightly slower compared to that observed with 50% priming cells (p=.0149; t test). We also validated the effectiveness of trans-killing with model antigens, thereby showing the robustness of trans-priming/killing (**Fig. S2 A-F)**. All of these *in vitro* killing studies show that trans-priming/killing effectively occurs, and that the EGFRvIII-primed circuit, thus, represents a promising strategy for preventing tumor escape due to heterogeneity.

We also hypothesized that T cells could also be locally primed by recognizing tissue-specific antigens (expressed on non-malignant cells) (**Fig. 1F**). For example, in the case of GBM, we might design T cell circuits that are primed by a brain-specific antigen, which then triggers local killing by inducing a CAR against the GBM antigens EphA2 and IL-13Rα2. A *brain antigen-primed* circuit might thereby provide a solution for treating EGFRvIII-negative GBM tumors. We bioinformatically identified two candidate brain surface proteins, Cadherin 10 (CDH10) – a brain-specific cadherin, and myelin oligodendrocyte glycoprotein (MOG) – a surface protein on the myelin sheath of neurons. The predicted tissue expression of these antigens is shown in **Fig. 1G and Fig. S7A**. For this study we focused primarily on using MOG as a priming antigen, as it proved to be more brain-specific (analysis of CDH10 as a priming antigen is shown in **Fig. S7A**). We identified antibodies that bind MOG and used them to construct cognate synNotch receptors that could be activated by cells expressing the mouse isoform of MOG (**Fig. 1H**), such that these receptors could be primed by endogenous mouse brain tissue. We then engineered CD8+ T cells with the **α-MOG synNotch** → **α-IL13Rα2/EphA2 CAR** circuit and co-cultured them with GBM target cells (here GBM6 PDX cell line), in the presence or absence of priming cells (L929 cells engineered to express MOG). We found that these T cells could effectively kill the GBM cells, but only in the presence of MOG+ priming cells (**Fig. 1I**). Importantly, these T cells did not show any killing of the priming cells. In summary, these *in vitro* studies show that there are multiple strategies for designing prime-and-kill circuits directed against GBM that execute trans-priming/killing, and thus have the potential to overcome antigen heterogeneity while still maintaining high specificity.

Based on these *in vitro* data, we next evaluated anti-tumor activities of these prime-and-kill CAR T cells in GBM xenograft mouse models. First, we wanted to confirm that the EGFRvIII-primed T cells could also perform trans-killing of EGFRvIII-GBM cells *in vivo*, but only in the presence of EGFRvIII+ priming cells. As proof of principle, we implanted NCG mice with dual tumors – in the brain we implanted a U87 tumor that was 50%EGFRvIII+ and 50% EGFRvIII-; in the flank we implanted a U87 tumor (EGFRvIII-only) (**Fig. 2A**). Here, the flank tumor represents a potentially cross-reactive normal tissue that expresses the killing antigens but not the priming antigen; in contrast, the brain tumor has both priming and killing antigens. On day 6 following the tumor inoculation, the mice received i.v. administration of prime-and-kill CAR T cells or control non-transduced T cells (n=6/group). All the mice treated with control T cells showed tumor growth at both sites and reached the euthanasia endpoint rapidly with a median survival of 25.5 days. The mice treated with prime-and-kill CAR T cells, in contrast, demonstrated significant suppression of the intracranial tumor growth compared with that in control mice (p< 0.001; t test). Importantly, however, the mice treated with the prime-and-kill CAR T cells did not show statistically significant suppression of the flank tumor compared with the control group (p=0.4; t test, **Fig. 2B**). The selective lack of killing in the non-priming flank tumor indicates that the cytotoxic activity of the prime-and-kill CAR T cells is spatially confined to the tumors expressing both priming and killing antigens.

**Figure 2.**
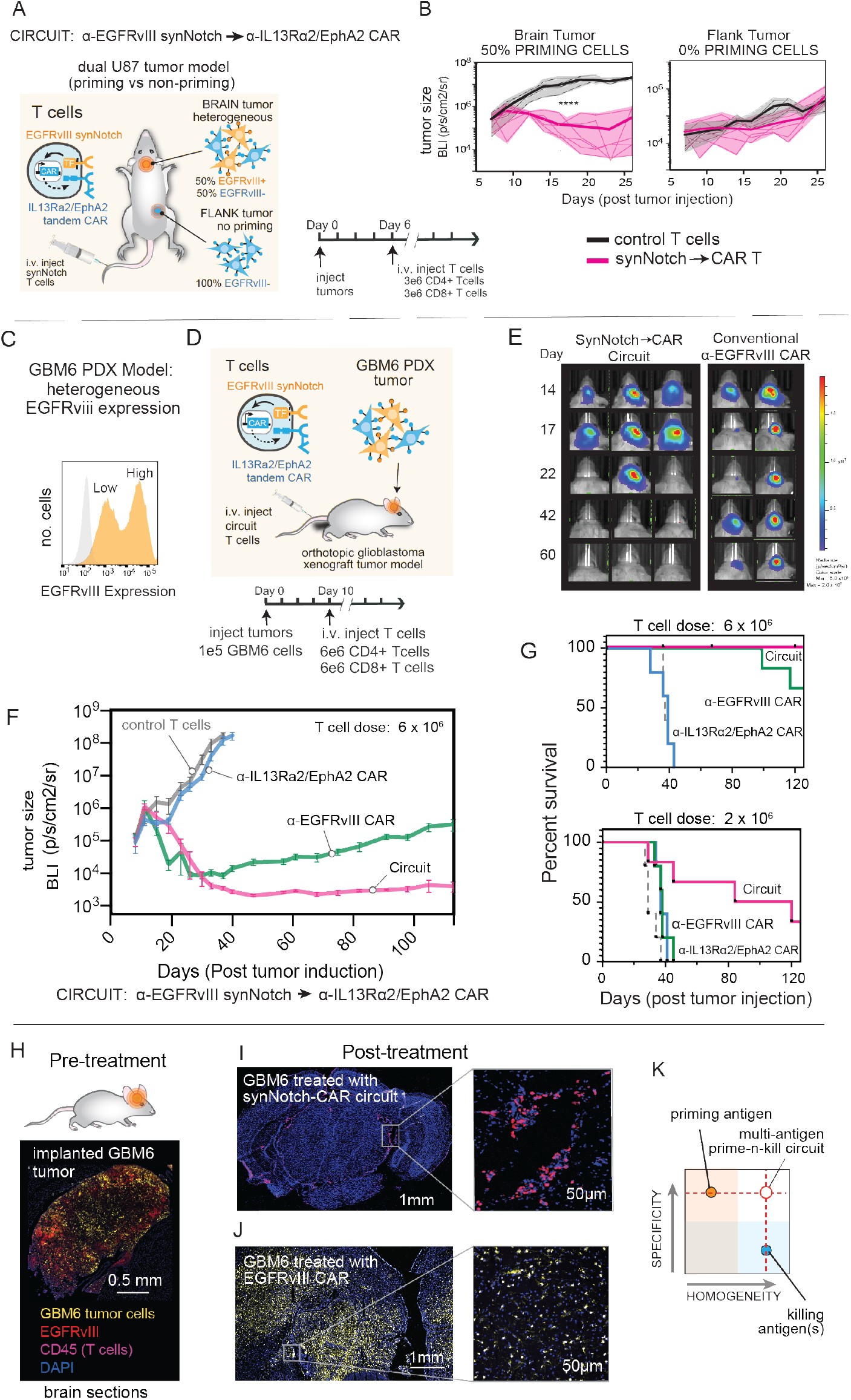
EGFRvIII neoantigen-primed T cell circuit shows improved specificity, efficacy and durability compared to individual parental constitutive CAR T cells in clearing heterogeneous GBM tumors. A. NCG mice were simultaneously implanted with two GBM tumors: i) heterogeneous tumor comprising EGFRvIII+ and EGFRvIII-U87 cells (1:1 ratio) in the brain, ii) homogeneous EGFRvIII-U87 cells in the flank (subcutaneous). Thus, both tumors express the killing antigens (EphA2 and IL13Rα2) but differ in expression of the EGFRvIII priming antigen. Mice were treated (6 days after tumor implantation) with intravenous infusion of 3 million of CD4+ and CD8+ prime-and-kill CAR T cells (n=6) or control non-transduced T cells (n= 6). B. Tumor size time course of two-tumor mouse experiment outlined in A, assayed by measuring luciferase luminescence. Tumor size curves for mice treated with prime-and-kill CAR T cells are shown in pink while curves for ones treated with non-transduced control T cells are shown in gray. Significant suppression in the size of brain tumor was observed in prime-and-kill CAR T cell treated mice (**** p < 0.00001; t test), while the flank tumor grew at the same rate as in the mice treated with non-transduced T cells. C. Patient derived xenograft (PDX) GBM model GBM6 shows intrinsic heterogeneity of EGFRvIII expression (low and high EGFRvIII population). FACS histogram showing anti-EGFRvIII staining. Further analysis of GBM6 cells is shown in **Fig. S4**. D. *In vivo* studies with GBM6 PDX tumors: NCG mice were orthotopically implanted in the brain with GBM6 PDX cells. Tumor cells were engineered to express mCherry and luciferase to allow for tracking of tumor size. Ten days following tumor implantation, the mice were infused intravenously with 3 million each of CD4+ and CD8+T cells. T cells expressed either: i) no construct (control) (n=5), ii) α-EGFRvIII synNotch→ α-IL13Rα2/EphA2 CAR circuit (n=6), iii) constitutively expressed α-EGFRvIII CAR (n=5), or iv) constitutively expressed α-IL13Rα2/EphA2 tandem CAR (n=5) E. Longitudinal bioluminescence imaging of GBM6 bearing mice treated with prime-and-kill CAR T cells and conventional α-EGFRvIII CAR T cells. Each column represents one mouse; each row represents time point of imaging. F. Time course tracking of tumor size determined by longitudinal bioluminescence imaging. Negative control treatment with non-transduced T cells is shown in black (n=5), prime-and-kill CAR circuit treatment is shown in pink (n=6), conventional α-EGFRvIII CAR treatment is shown in green line (n=6), and constitutive α-IL13Rα2/EphA2 tandem CAR in blue (n=5). Mice treated with constitutive α-IL13Rα2/EphA2 tandem CAR failed to control tumor growth. Treatment with conventional α-EGFRvIII CAR T cells resulted in early tumor regression followed by recurrence in all mice (n=6). In contrast, all mice treated with the prime-and-kill CAR T cells showed complete clearance of tumor (p< 0.0001 t-test untransduced vs prime-and-kill CAR T cells). Error bars represent mean ± SEM of 5-6 individual mice from one experiment. An independent replicate of this experiment is shown in **Fig S3**.**C**. G. Kaplan-Meier survival curves over 100 days for high dose (6 × 10^6^ cells, upper panel) and low dose (2 × 10^6^ cells, lower panel). Treatment with constitutive α-IL13Rα2/EphA2 tandem CAR did not confer survival benefit (median survival, 39d for α-IL13Rα2/EphA2 CAR versus 37d for untransduced T cells). At low dose, all the mice treated with constitutive EGFRvIII CAR succumbed to tumor-induced death (median survival of 40d) while high dose treatment resulted in 2 out of 6 mice dead by day 117. In contrast, all mice treated with the prime-and-kill CAR T cells (high dose) showed complete clearance of tumor and mice survived for more than 125 days, except for three mice that were euthanized because of unrelated infection. Treatment of mice with low dose of prime-and-kill CAR T cells resulted in survival of 3 out of 6 mice for over 100 days. Statistical significance was calculated using the log-rank Mantel–Cox test. H. Representative fluorescence microscopy of GBM6 xenograft tumor (prior to T cell treatment) shows heterogeneous expression of EGFRvIII (red) (tumor in yellow). Images are taken 10 days post tumor implantation. I. Fluorescent microscopy reveals clearance of engrafted GBM6 xenograft (lack of mCherry tumor cells (yellow) following treatment with synNotch-CAR circuit T cells. Retention of prime-and-kill CAR T cells (stained for human CD45, in red) is observed in the brain parenchyma and meninges. J. Representative image of tumor recurrence (mCherry positive tumor cells, false-colored in yellow) with loss of EGFRvIII expression (in red) for xenograft tumors treated with conventional α-EGFRvIII CAR T cells. K. T cell circuits can be designed to combine recognition of two imperfect but complementary antigens: a highly specific but homogeneous as a priming antigen, and a non-specific but homogenous set of antigens as killing targets. By applying a specificity pre-filter (requirement for priming antigen) over a less constrained killing mechanism, these combinatorial antigen recognition circuits may enable the engineered T cells to overcome tumor antigen heterogeneity while also eliminating toxicity caused by insufficient specificity (on-target, off-tumor attack of normal tissues).

We also performed a systematic comparison of killing of implanted tumors with 0%, 50%, and 100% EGFRvIII+ U87 cells (**Fig**.**S3 A, B)**. We found that prime-and-kill CAR T cells did not show any clearance of the 0% EGFRvIII-positive tumors, but showed equally effective tumor clearance of both the 50% and 100% EGFRvIII-positive tumors (controls did not show clearance). Thus, in this context, prime-and-kill CAR T cells can recognize and effectively overcome tumors with heterogeneous EGFRvIII expression, but do so in a specific, priming antigen-gated manner.

We then sought to evaluate the efficacy of prime-and-kill CAR T cells in a tumor model which exhibits naturally occurring heterogeneity of EGFRvIII expression. We identified the GBM6 patient-derived xenograft (PDX) tumor as an aggressive GBM model that shows intrinsic EGFRvIII heterogeneity (*23*) (**Fig. 2C**). Equally importantly, GBM6 tumors show a highly reproducible ability to evade treatment by the EGFRvIII single antigen CAR *in vivo* (**Fig. 2F**). When mice are treated with the α-EGFRvIII CAR, the GBM6 tumors shrink dramatically, but then slowly and steadily recur with high reproducibility (**Fig. 2F, Fig. S3C**, discussed below). These recurrent tumors show loss of EGFRvIII expression (**Fig. 2J**) (we confirmed *in vitro* that there is a subpopulation of GBM6 cells that show undetectable EGFRvIII antigen and are resistant to killing by the conventional EGFRvIII CAR – **Fig. S4 C**,**D**). Thus, GBM6 tumors mimic the heterogeneity-based escape observed in α-EGFRvIII CAR clinical trials, and therefore represents an ideal tumor model in which to evaluate alternative circuits that could overcome these problems.

We tested treatment of NCG mice bearing GBM6 tumors in the brain with T cells bearing our EGFRvIII-primed circuit, non-transduced (control) T cells or T cells constitutively expressing either the α-EGFRvIII CAR or α-IL13Rα2/EphA2 tandem CAR (**Fig. 2 F,G**). All of the mice receiving the non-transduced control T cells (n=5) died of tumor progression by day 43 post-tumor inoculation (**Fig. 2 E, F**). Treatment with the constitutive α-IL13Rα2/EphA2 tandem CAR was largely ineffective (despite effective cytotoxicity of these T cells *in vitro*) (**Fig. S4D**). Treatment with α-EGFRvIII CAR T cells yielded initial tumor shrinkage, but consistently (n=6) resulted in recurrence of EGFRvIII-negative tumors in all mice (3 of 6 mice died of tumor progression by Day 125) (**Fig. 2 E, F, J**). In striking contrast, all of the mice (n=6) treated with the prime- and-kill CAR T cells showed long-term complete remission of the GBM6 tumors. This more durable and complete tumor clearance was highly reproducible (**Fig. S3C**), and was also reflected in significantly increased survival rates of mice treated with the prime-and-kill circuit (performed at multiple doses) (**Fig. 2G**).

We performed post-mortem immunofluorescence analysis of mice treated with either the EGFRvIII CAR or the prime-and-kill circuit. The brains of mice treated with the prime-and-kill circuit showed an absence of GBM6 tumor cells (consistent with tumor clearance), but showed persistence of CAR T cells (pink) in the brain parenchyma and meninges (**Fig. 2I**). In contrast, the constitutive EGFRvIII CAR-treated brains showed extensive GBM6 tumor cells (yellow – consistent with recurrence), but a loss of EGFRvIII antigen (red) and no surviving CAR T cells (**Fig. 2J**). In summary, the *in vivo* tumor killing studies support the concept that a dual-antigen circuit can combine the specificity of a priming antigen with the homogeneity of a killing antigen to achieve far more specific but complete killing than is possible with CARs targeted for either single antigen.

To allow for direct visualization of T cell priming *in vivo*, we fused a GFP tag onto the induced α-IL13Rα2/EphA2 CAR. Six days after T cell infusion, analysis of brain slices from the recipient mice revealed GFP+ prime-and-kill CAR T cells (co-stained for human CD45) in the tumor (**Fig 3A**). In contrast, no primed T cells were found outside of the tumor (adjacent brain tissue), nor in the spleen (**Fig 3A**). Furthermore, we performed intravital imaging of these prime-and-kill T cells two days after injection, which showed a large number of primed (green) T cells in the tumor, as well as T cells that became green as they approached the tumor (**Fig. 3B, Movie S2**). Significantly, these movies show that the primed T cells remain stably localized within the tumor (presumably interacting with target cells), and do not rapidly traffic in or out of the tumor. This behavior may help explain the highly specific and localized killing by these T cells.

**Figure 3.**
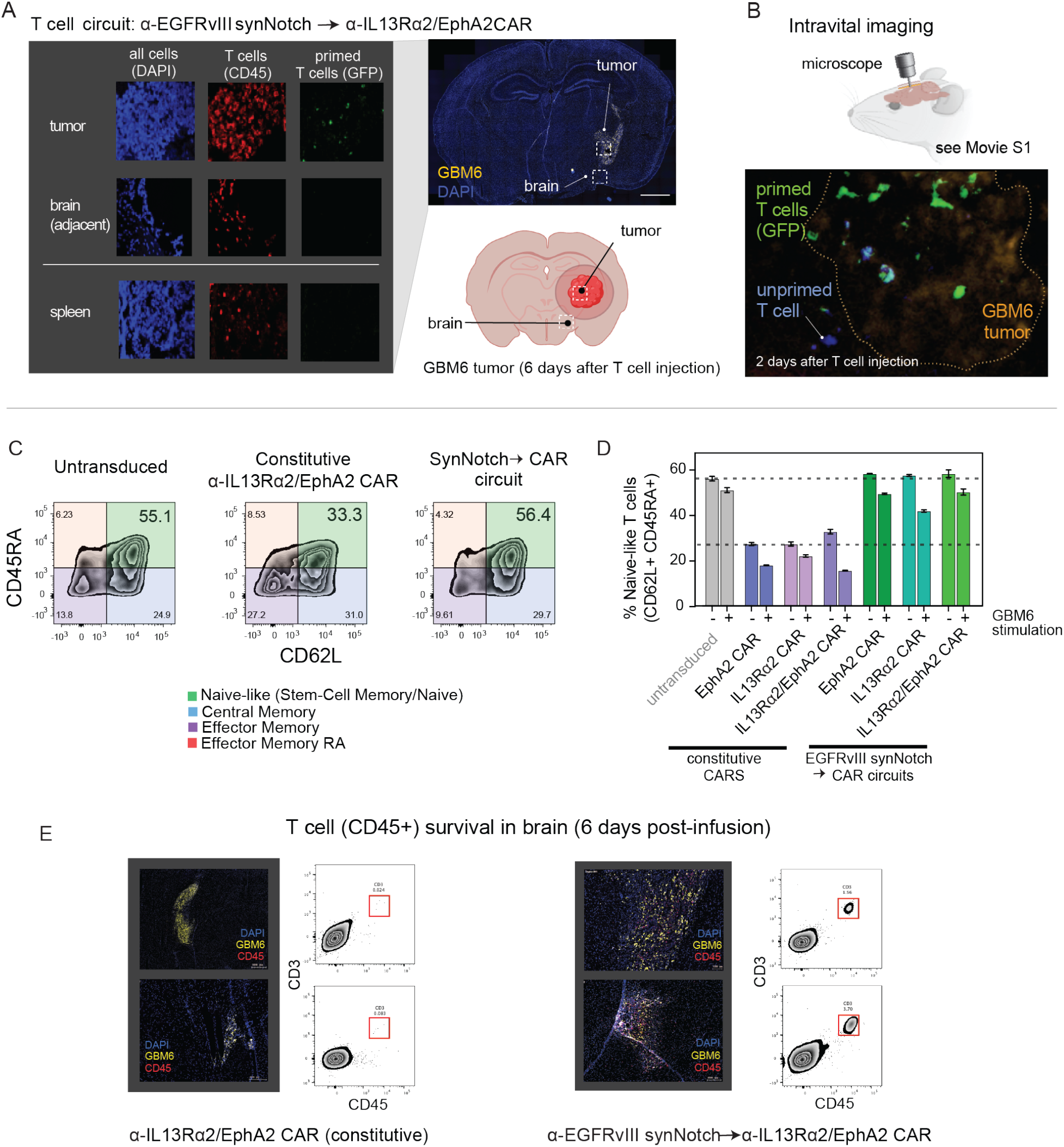
EGFRvIII SynNotch → CAR circuit T cells show tumor localized priming and maintain larger fraction of naïve-state cells compared to standard constitutive CARs. A. GBM6 tumor-bearing mice were euthanized six days after infusion with EGFRvIII SynNotch →CAR-GFP circuit T cells. Representative confocal fluorescent microscopy of brain slices reveal primed GFP+ T cells (hCD45 stain in red) in the tumor bed (yellow) but not in the periphery. Circuit CAR T cells express GFP upon priming. Circuit CAR T cells (red) in the spleen do not express GFP. Insets of single stained images are enlargements of outlined regions in the main image. DAPI stain was used for nuclear segmentation. CD45 (AF647) and GFP were then co-localized with the nuclear stain to quantify T cells (CD45+) and the fraction that were primed (GFP+). Images are representative from 1 of 3 mice. B. Intravital imaging (Two-photon *in vivo* laser scanning) of circuit CAR T cells show dynamic priming within the GBM6 xenograft tumor. Tumors were implanted at a depth of 3mm below the right frontal cortex and cranial windows were implanted; tumor cells, orange; unprimed circuit CAR T cell, blue; primed circuit CAR T cell, green. See Supplemental Movie S2. C. Surface expression of CD45RA and CD62L to distinguish naïve-like cells (CD45RA+CD62L+), central memory cells (CD45RA−CD62L+), effector memory cells (CD45RA−CD62L−), and effector memory RA cells (CD45RA+CD62L-) (representative of 3 experiments). Comparison on cell phenotype distribution for untransduced T cells, T cells with constitutive α-IL13Rα2/EphA2 tandem CAR, and T cells with synNotch→CAR circuit. Percentage of cells in the naïve/memory stem cell state is highlighted. D. Percentage of CD62L+CD45RA+ phenotype cells in all relevant CAR T cells for this study, including constitutive α-EphA2 CAR, constitutive α-IL13Rα2 CAR, and constitutive α-IL13Rα2/EphA2 tandem CAR, as well as the EGFRvIII synNotch-primed versions of these circuits. Data are shown with and without GBM6 cells (after 24 hours). Bars show the percent of T cell population with naïve-like T cells (n = 3). See **Fig. S5** for further phenotypic analysis of this and other engineered T cells from this study. E. SynNotch→ CAR circuit T cells show improved persistence *in vivo* compared to constitutive CAR T cells. GBM6 tumor-bearing mice were euthanized six days after infusion of either the constitutive tandem CAR T cells (left panels) or the EGFRvIII prime-and-kill circuit T cells (right panels). Two representative confocal fluorescent microscopy of brain slices reveal few detectable constitutive CAR T cells (hCD45 stain in red) in the tumor (yellow). Single-cell suspensions of tumor (from day 6 after T cell infusion) were labelled with CD3 and CD45 for analysis by flow cytometry, showing few T cells (CD3+CD45+). In Right panel: On day 6, mice were euthanized and tumor tissue was collected and dissociated. Single-cell suspensions were labelled with CD3 and CD45 for analysis by flow cytometry. In contrast brain slices derived from mice treated with EGFRvIII SynNotch →CAR circuit T cells reveal high number of T cells (hCD45 stain in red) in the tumor (yellow). Single-cell suspensions (day 6) were also show far higher number of the circuit T cells by flow cytometry.

One the most surprising findings of these *in vivo* studies was the significantly better tumor clearance capacity of the prime-and-kill T cells compared to the constitutive α-IL13Rα2/EphA2 CAR T cells, as both sets of T cells kill using the same CAR molecule, and both are equally effective at tumor killing *in vitro* (**Fig. S4D**). These observations suggested that there were additional features of synNotch-induced CAR circuits that yielded significantly improved anti-tumor activity *in vivo*. A general challenge in treating solid cancers with CAR T cells is exhaustion of the T cells, which prevents durable anti-tumor activity. Recent studies have indicated that tonic signaling by constitutively expressed CARs can play a significant role in increasing their susceptibility to exhaustion (*24, 25*). We therefore examined the differentiation state of the different types of T cells by flow analysis, and found that all of the synNotch→CAR T cells used in this study showed significantly higher fraction of cells in a naïve-like state (CD62L+ CD45RA+, naïve or stem central memory) compared to the equivalent constitutive CAR T cells (**Fig. 3C,D**). Furthermore, when we directly investigated T cell persistence in vivo, 6 days after infusion into the mouse, we observed a high number of the synNotch→CAR circuit T cells. In contrast, we found no surviving constitutive tandem CAR T cells by this time (**Fig. 3E**). Together, these findings are consistent with a simple model: synNotch→CAR circuits prevent the tonic signaling normally observed in constitutively expressed CARs, thereby allowing the T cells to maintain a more naïve-like state that is less prone to exhaustion. Thus, local, transient priming of CAR expression not only increases targeting specificity, but also appears to yield a more potent and persistent T cell state.

We also wanted to test whether brain-specific antigen-primed T cells could be effective *in vivo*. Thus, we treated NCG mice implanted with intracranial GBM6 PDX tumors with T cells bearing the **α-MOG synNotch**→ **α-IL13Rα2/EphA2 CAR** circuit (**Fig. 4A**). The GBM6 tumor cells do not express MOG, therefore in order for the T cells to be primed, they must be primed by MOG endogenously expressed in the host mouse brain (**Fig. 1I**). We found that the MOG-primed T cells were highly effective at clearing the GBM6 tumor and increasing mouse survival (**Fig. 4 B, C**). In mice in which GBM6 tumors are implanted in the flank instead of the brain, tumors are not cleared by the MOG-primed T cells (**Fig. S7F**), consistent with a need for a localized brain signal to license the T cells for killing. Mice treated with the **α-CDH10 synNotch**→ **α-IL13Rα2/EphA2 CAR** circuit also showed effective killing of GBM6, but in this case showed poor discrimination in killing of brain vs flank tumors, suggesting that MOG is superior to CDH10 as a tightly restricted brain-specific priming antigen (**Fig. S7F**).

**Figure 4.**
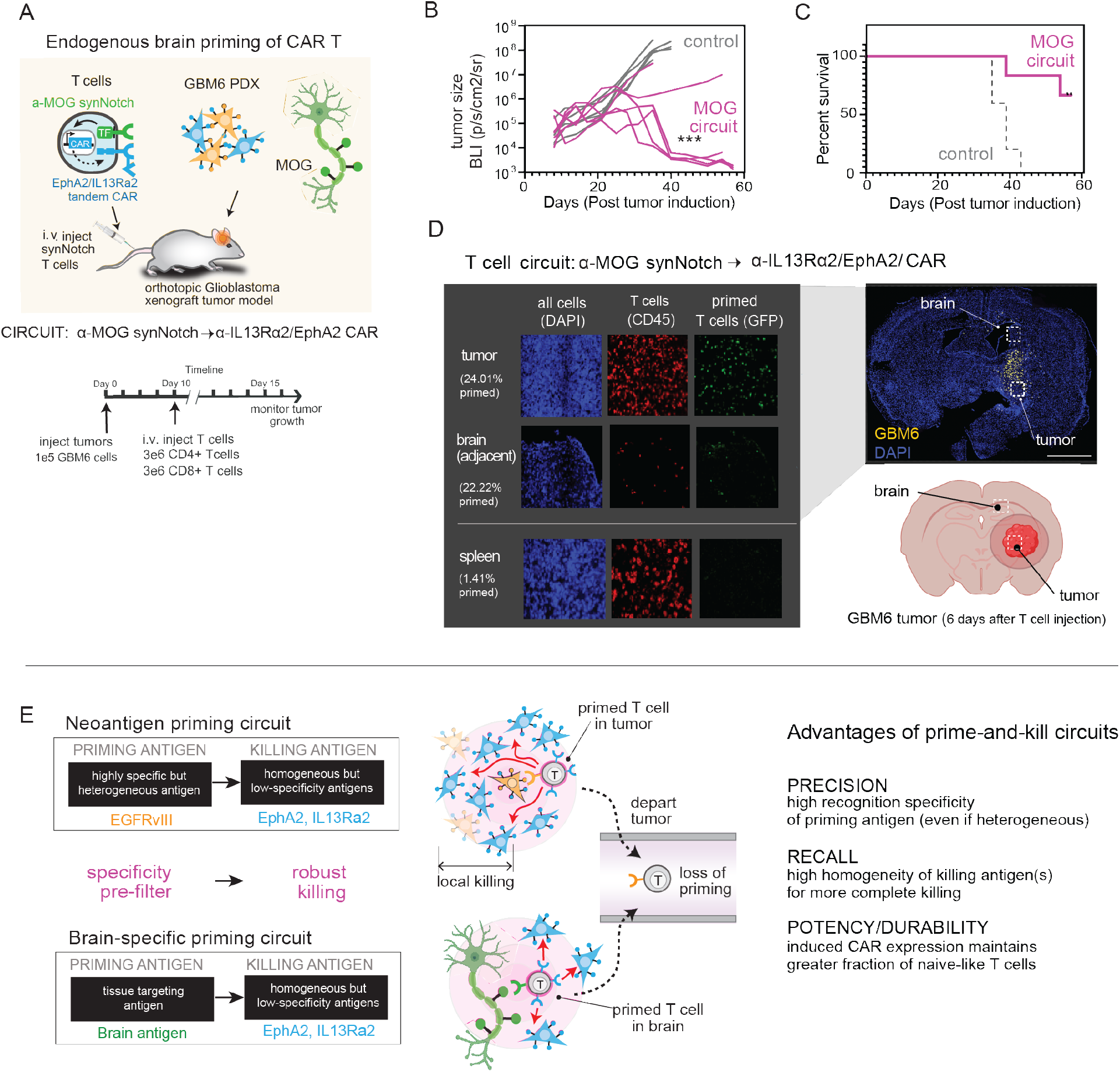
Tissue-specific priming of SynNotch CAR circuit T cell (by MOG antigen) induces killing of GBM6 brain tumors. A. NCG mice were stereotactically implanted in the brain with GBM6 PDX cells. Tumor cells were engineered to express mCherry and luciferase to allow for tracking of tumor size. Ten days following tumor implantation, mice were infused intravenously with 3 million each of CD4+ and CD8+T cells. T cells expressed either: i) no construct (untransduced control) (n=5), or ii) α-MOG synNotch→ α-IL13Rα2/EphA2 CAR circuit (n=6). B,C. Tumor size and survival monitored over time. Tumor size was determined by longitudinal bioluminescence imaging. Negative control treatment (untransduced T cells) is shown in grey, MOG SynNotch→CAR circuit treatment is shown in pink. Compared to non-transduced treatment group, mice treated with the α-MOG-prime-and-kill CAR T cells (4 out of 6 mice) showed strong anti-tumor response (p<0.001 by t test with Holm-Sidak correction for multiple comparisons) and survived for 60 days [p = 0.05 Log-rank (Mantel-Cox) test]. D. Visualizing T cell priming in brain and tumor. GBM6 tumor-bearing mice were euthanized six days after MOG SynNotch →CAR circuit T cell infusion. Representative confocal fluorescent microscopy of sections obtained from circuit T cell-treated mice reveals primed T cells (GFP+ - priming; red - T cells identified with human CD45 stain) in the tumor (yellow) and in the infiltrative parts of the tumor. Brain tissue adjacent to the tumor shows a lower overall number of T cells, but a similar percentage are primed (GFP+). Circuit CAR T cells (red) in the spleen do not show any significant priming. Insets (single stained images) are enlargements of outlined regions in the main image. E. synNotch→CAR circuits combine advantageous features of multiple individually imperfect antigens to achieve far better therapeutic tumor recognition behavior. A priming antigen (either neoepitope or tissue-specific antigen) can be used to achieve high precision killing, even if the priming antigen is heterogeneously expressed or expressed on non-cancer cells. A homogeneous killing antigen (or combination of antigens) can then be subsequently targeted to achieve complete tumor killing (e.g. high recall). In addition, we find that synNotch→CAR T cells are more naive-like and more durable than their equivalent constitutively expressed CAR T cells.

Post-mortem immunofluorescence analysis of mice treated with the MOG-primed T cells revealed a high number of T cells in the tumor, with many of these in a primed state (visualized by GFP-fused CAR) (**Fig. 4D**). In the tumor-adjacent brain tissue we observed fewer T cells, but these were also primed (GFP+) (no primed T cells were observed in the spleen). These results are consistent with T cell priming throughout the brain, presumably combined with more significant T cell expansion in the tumor, resulting from CAR activation and release of proliferative cytokines.

The effectiveness of this brain-specific antigen-primed circuit represents a novel advance in engineering therapeutic cells – not only does this circuit represent a possible way to treat EGFRvIII^-^ GBM, but it also may provide a general strategy for engineering brain-targeted cell therapies to treat a broad range of neurological diseases, including other brain tumors, neuroinflammation or neurodegeneration.

Together, these results show that that there are multiple ways to engineer synNotch→CAR circuits that effectively combine recognition of complementary antigens that are imperfect as single antigen targets. One can build circuits that prime based on a highly specific neoantigen, like EGFRvIII, or alternatively a tissue-specific antigen like MOG. Critically, these priming antigens need not be on all tumor cells, or on any of the tumors cells at all (in the case of tissue-specific priming). Once primed, the T cells can then be programmed to execute complete tumor killing by targeting CAR antigens that are homogeneous on the tumor, even if they have imperfect specificity by themselves. By integrating information from multiple antigens and multiple cells, these circuits essentially give us improved capability for nuanced recognition of a tumor as a complex tissue, and thus open up many new possibilities for how to recognize and attack tumors in safer and more specific ways. Other related strategies combine CARs and bispecific engagers to integrate multi-antigen combinations (*26*).

We show here that synNotch→CAR T cells have multiple features distinguishing them from conventional CAR T cells, which may prove to be highly advantageous in treating solid cancers such as GBM. First, their multiantigen recognition increases discrimination between tumor and normal cells. Second, the ability of these circuits to mediate trans-priming/killing allows them to overcome escape via target antigen heterogeneity. Finally, the simple act of placing CAR expression under regulated control appears to maintain the T cells in a far more naïve-like state that is more durable and less subject to exhaustion. Similar improved anti-tumor activity has been observed for synNotch→ CAR T cells targeting other cancers beyond GBM (companion submitted manuscript – 27), suggesting that these circuits may provide a highly effective general strategy for treating many solid cancers.

## NOTES

## ACKNOWLEDGEMENTS

We thank J. Williams, R. Hernandez-Lopez, V. Tieu, N. Blizzard, J. Shon, M. Shahin and members of the W.A.L. lab and H.O. lab for technical assistance, advice and helpful discussions. This work was supported by NIH grants P50GM081879 (W.A.L), R01 CA196277 (W.A.L), Howard Hughes Medical Institute (W.A.L), 1R35 NS105068 (H.O.), UCSF Glioblastoma Precision Medicine Program (W.A.L and H.O.) and Parker Institute for Cancer Immunotherapy (W.A.L. and H.O.).

## AUTHOR CONTRIBUTIONS

Conceptualization, J.H.C., P.B.W., K.T.R., H.O., and W.A.L.; Brain-Priming Circuits, M.S. and J.H.C.; Formal Analysis, J.H.C. and P.B.W.; Funding Acquisition, H.O. and W.A.L.; Investigation, J.H.C., P.B.W., M.S., R.D.G., D.A.C, W.Y., K.M.D., N.A.K., A.W.L, R.D., J.C., and J.D.B; Methodology, J.H.C. and P.B.W.; Project Administration, J.H.C., P.B.W., K.T.R., H.O., and W.A.L. Resources, S.S.; Supervision, K.T.R., H.O., and W.A.L.; Visualization, J.H.C., P.B.W., and W.A.L.; Writing – original draft, J.H.C., P.B.W., K.T.R., H.O., and W.A.L.; Writing – review and editing, J.H.C., P.B.W., R.D.G., K.T.R., H.O., and W.A.L.

## SUPPLEMENTARY MATERIALS

Materials and Methods

Figs. S1-S7

Movies. S1-2

