## Supplementary Data for "Multi-antigen recognition circuits overcome challenges of specificity, heterogeneity, and durability in T cell therapy for glioblastoma"

**This PDF file includes:**

Materials and Methods
Figs. S1 to S7
Captions for Movies S1 to S2

**Other Supplementary Materials for this manuscript include the following:**

Movies S1 to S2

**Materials and Methods**

CONTACT FOR REAGENT AND RESOURCE SHARING

Reagent requests should be directed and will be fulfilled by lead author Wendell Lim. To ensure a fast response, please copy Noleine Blizzard and Michael Broeker in any requests related to the paper.

Source of Primary Human T cells

Blood was obtained from Blood Centers of the Pacific (San Francisco, CA) as approved by the University Institutional Review Board. Primary CD4+ and CD8+ T cells were isolated from anonymous donor blood after apheresis (described in METHOD DETAILS).

Construct Design

SynNotch receptors were built by fusing the EGFRvIII 139 scFv (*1*), MOG M26 scFv (*2*), and CDH10 (gift from Sidhu lab) to the mouse Notch1 (NM_008714) minimal regulatory region (Ile1427 to Arg1752) and Gal4 DBD VP64. All synNotch receptors contain an n-terminal CD8a signal peptide (MALPVTALLLPLALLL HAARP) for membrane targeting and a myc-tag (EQKLISEEDL) or flag-tag (DYKDDDDK) for easy determination of surface expression with a-myc A647 (cell-signaling #2233) or a-flag A647 (RND systems #IC8529R); see Morsut et al. (*3*) for receptor synNotch receptor peptide sequences). The receptors were cloned into a modified pHR'SIN:CSW vector containing a PGK or SFFV promoter for all primary T cell experiments. The pHR'SIN:CSW vector was also modified to make the response element plasmids. Five copies of the Gal4 DNA binding domain target sequence (GGAGCACTGTCCTCC GAACG) were cloned to a minimal CMV promoter. Also included in the response element plasmids is a PGK promoter that constitutively drives mCherry or BFP expression to easily identify transduced T cells. Inducible CARs were built by fusing EphA2 scFv (*4*), IL13 Mutein [E13K,K105R] (*5*), or IL13 Mutein [E13K,K105R]-G4Sx4-EphA2 scFv (*4, 5*) to the hinge region of the human CD8α chain and transmembrane and cytoplasmic regions of the human 4-1BB, and CD3z signaling endodomains. The inducible CAR constructs were cloned via a BamHI site in the multiple cloning site 3' to the Gal4 response elements. For some inducible CAR vectors, the CARs were tagged c-terminally with GFP/BFP or contain myc/flag tag to verify surface expression. All constructs were cloned via in-fusion cloning (Clontech #ST0345).

Primary Human T Cell Isolation and Culture

Primary CD4+ and CD8+ T cells were isolated from anonymous donor blood after apheresis by negative selection (STEMCELL Technologies #15062 and #15063). Blood was obtained from Blood Centers of the Pacific, as approved by the University Institutional Review Board. T cells were cryopreserved in RPMI-1640 (UCSF cell culture core) with 20% human AB serum (Valley Biomedical, #HP1022) and 10% DMSO. After thawing, T cells were cultured in human T cell medium consisting of X-VIVO 15 (Lonza #04-418Q), 5% Human AB serum, and 10 mM neutralized N-acetyl L-Cysteine (Sigma-Aldrich #A9165) supplemented with 30 units/mL IL-2 (NCI BRB Preclinical Repository) for all experiments except for the IncuCyte experiments. IncuCyte experiments were cultured in RPMI-1640 (UCSF cell culture core) with 5% human AB serum (Valley Biomedical, #HP1022) supplemented with 30 units/mL IL-2 (NCI BRB Preclinical Repository).

Lentiviral Transduction of Human T Cells

Pantropic VSV-G pseudotyped lentivirus was produced via transfection of Lenti-X 293T cells (Clontech #11131D) with a pHR'SIN:CSW transgene expression vector and the viral packaging plasmids pCMVdR8.91 and pMD2.G using Fugene HD (Promega #E2312). Primary T cells were thawed the same day and, after 24 hr. in culture, were stimulated with Human T-Activator CD3/CD28 Dynabeads (Life Technologies #11131D) at a 1:3 cell:bead ratio. At 48 hr., viral supernatant was harvested and in some assays concentrated using Lenti-X concentrator (Clonetech #631231). The primary T cells were exposed to the virus for 24 hr. At day 4 after T cell stimulation, the Dynabeads were removed, and the T cells were expanded until day 9 when they were rested and could be used in assays. T cells were sorted for assays with a Beckton Dickinson (BD) FACs ARIA Fusion or Sony SH800S Cell Sorter. AND-gate T cells exhibiting basal CAR expression were gated out during sorting.

Cancer Cell Lines

The cancer cell lines used were K562 myelogenous leukemia cells (ATCC #CCL-243), L929 mouse fibroblast cells (ATCC# CCL-1), U87 MG GBM cells (ATCC #HTB-14), and GBM6 PDX cells (generous gifts from Dr. Frank Furnari at Ludwig Institute and UCSD). U87-EGFRvIII-negativeluciferase (Ohno et al., 2013) and U87 MG were lentivirally transduced to stably express GFP or mCherry, respectively, under control of the spleen focus-forming virus (SFFV) promoter. At 72 hours after transductions, cells were sorted on an Aria Fusion cell sorter (BD Biosciences) on the basis of GFP expression to be 100% GFP or mCherry positive and subsequently expanded. All cell lines were sorted for expression of the transgenes. U87-luciferase and U87-luciferase-mCherry cells were stably transduced with non-mutated EGFR using a retroviral construct (a gift from Matthew Meyerson; Addgene plasmid # 11011) to generate EGFRvIII-negative cell lines which grow at a similar rate as the EGFRvIII-positive U87 cell line (see Fig. S2). Overexpression of wild-type EGFR is relevant to human GBM with EGFR amplification (*6*). GBM6 were lentivirally transduced to stably express both mCherry and firefly luciferase. These cells were cultured in DMEM F12 media, with supplements of EGF (20 μg/mL), FGF (20 μg/mL), and heparin (5 μg/mL). K562s were lentivirally transduced to stably express surface CDH10 (CDH10 extracellular membrane was fused to the PDGF transmembrane domain). K562s and L929 were lentivirally transduced to stably express full length MOG.

In Vitro Stimulation of SynNotch T cells

For all in vitro synNotch T cell stimulations co-cultured with U87, 1 x $10^4$ U87s were cultured overnight in a flat bottom 96-well tissue culture plate. Next morning, 1 x $10^4$ - 5 x $10^4$ T cells were added to the flat bottom 96-well tissue culture plate and the co-cultures were analyzed at 24-96 hr. for activation and specific lysis of target tumor cells. For all in vitro synNotch T cell stimulations co-cultured with GBM6 and T cells, 1 x $10^4$ GBM6s were cultured overnight in a flat bottom 96-well tissue culture plate. Next morning, 1 x $10^4$ T cells were added to the flat bottom 96-well tissue culture plate and the co-cultures were analyzed at 24-96 hr. for activation and specific lysis of target tumor cells. For all in vitro synNotch T cell stimulations co-cultured with three different cell populations, the target cells (GBM6) were cultured at 1 x $10^4$ cells and priming cells (either K562 or L929) were cultured at 1 x $10^4$ cells overnight in a flat bottom 96-well tissue culture plate. Next morning, 1 x $10^4$ T cells were added to the flat bottom 96-well tissue culture plate and the co-cultures were analyzed at 24-96 hr. for activation and specific lysis of target tumor cells. All flow cytometry was performed using BD LSR II or Attune NxT Flow Cytometer and the analysis was performed in FlowJo software (TreeStar).

Assessment of SynNotch AND-Gate T Cell Cytotoxicity

CD8+ synNotch AND-Gate T cells were stimulated for 24-96 hr. as described above with target cells expressing the indicated antigens. The level of specific lysis of target cancer cells was determined by comparing the fraction of target cells alive in the culture compared to treatment with non-transduced T cell controls unless stated otherwise. Cell death was monitored by shifting of the target cells out of the side scatter and forward scatter region normally populated by the target cells. Alternatively, cell viability was analyzed using the IncuCyte Zoom system (Essen Bioscience). Tumor cells were plated into a 96-well plate at a density of $1.0 \times 10^4$ cells per well in triplicate overnight. T cells were added into each well next day at a final volume of 200 µl per well. The target cells and T cells were co-cultured as described above. 2 fields of view were taken per well every 15 minutes. The mean florescence intensity (MFI) was calculated using IncuCyte Zoom software (Essen BioScience) in order to determine the target cell survival. The data were summarized as mean ± SEM.

Statistical Analysis and Curve Fitting

Statistical significance was determined by specific tests and presented as means ± standard error mean (SEM) or means ± standard deviations (SD) as indicated in the figure legends. The Kaplan-Meier estimator was used to generate survival curves, and differences in survival distributions were assessed using Log-Rank test. All p values are provided in the figures or their legends. All statistical analyses were performed with Prism software version 7.0 (GraphPad).

In vivo Mouse Experiments

All mouse experiments were conducted according to Institutional Animal Care and Use Committee (IACUC)–approved protocols. For orthotopic heterogeneous model with U87, a mixture of $1.5 \times 10^4$ U87-luc-mCherry and $1.5 \times 10^4$ U87-luc-EGFRvIII-negativeGFP cells was implanted intracranially into 6- to 8-week-old female NCG mice (Charles River), with 6-10 mice per group. For homogeneous U87-luc-GFP-EGFRvIII-positive model, $3 \times 10^4$ cells were injected into the brains of NCG mice. For orthotopic heterogeneous model with GBM6, $1.0 \times 10^5$ GBM6-luc-mcherry cells were implanted intracranially into 6- to 8-week-old female NCG mice with 5-10 mice per group. A stereotactic surgery for tumor cell inoculation was performed with the coordination of the injection site at 2 mm right and 1 mm anterior to the bregma and 3 mm into the brain. Before surgery and for 3 days after surgery, mice were treated with an analgesic and monitored for adverse symptoms in accordance with the IACUC. In subcutaneous model, NCG mice were injected with either $1.0 \times 10^6$ U87-Luc-mcherry+ or $1.2 \times 10^5$ GBM6-luc-mcherry cells subcutaneously in 100 µl of HBSS on day 0. Tumor progression was evaluated by luminescence emission on a Xenogen IVIS Spectrum after intraperitoneal D-luciferin injection according to the manufacturer's directions (GoldBio). Prior to the treatment, mice were randomized such that initial tumor burden in the control and treatment groups were equivalent. Mice were treated with $6.0 \times 10^6$ engineered or the matched number of non-transduced T cells intravenously via tail vein in 100 µl of PBS. Survival was evaluated over time until predetermined IACUC-approved endpoint (hunching, neurological impairments such as circling, ataxia, paralysis, limping, head tilt, balance problems, seizures) was reached (n = 6 to 10 mice per group).

Two-Photon In Vivo Microscopy

Intravital two-photon images were acquired with a Zeiss LSM 780 NLO equipped with a Ti:Sapphire laser (MaiTai HP, Spectra Physics) tuned to 760 nm (for excitation of tagBFP+ synNotch CAR T cells and mCherry+ tumor) and 900 nm (for excitation of GFP+ synNotch CAR T cells), respectively, and focused through a Zeiss 20× water immersion objective (numeric aperture of 1.0). Before imaging, mice were anesthetized with isoflurane and the head plate was fixed into the head posts of a custom-made moving stage (Thorlabs, UC Berkeley Physics Machine Shop). Anesthesia was maintained at 1% isoflurane through a nose cone, and body temperature was kept stable via a temperature-controlled heating pad. Images of 598 × 598 µm2 areas of in vivo tumors were acquired at 512 x 512 pixel resolution for standard images and 1,024 × 1,024 pixel resolution for higher resolution images. Volume images were acquired over a 30- to 200-µm Z range in 5- or 10-µm steps. Time-lapse datasets were acquired either in single planes over time periods up to 45 min, or in combined time + Z series over a 598×598 area (X×Y) with variable Z ranges (Z= 5- to 100um) with 1- to 5um steps. Movies were processed in Zen software; for 3D reconstructions.

Cranial Window Implantation

8-week-old NCG mice underwent tumor implantation with GBM6 xenograft as described above. 10-12 days post-tumor injection, mice underwent the implantation of a cranial window and custom designed titanium head plate (UC Berkeley Physics Machine Shop; design by Kira Poskanzer, PhD) according to the following procedure. The cranial window consisted of a No.1 4mm glass coverslip (Warner Scientific) glued onto a No. 1 3mm glass coverslip with optical adhesive (Norland 71, Norland Products) and cured with 365nm UV light for 15 minutes. Five hours before surgery, mice were injected once intraperitoneally with dexamethasone (2.8mg/kg). At time of surgery, mice were anesthetized with 1.5-3% isofluorane and secured into a stereotactic frame with ear bars. The head was shaved, an ovular flap of skin was removed from the top of the head and skin margins were glued down with bio-compatible cyanoacrylate glue (VetBond, 3M). The titanium headplate was superglued (KrazyGlue) to the skull and secured with dental cement at the end of the surgery (C&B Metabond Kit, Parkell). A 3mm diameter craniotomy was performed using a Foredom micro-drill with 1mm and 0.5mm carbide drill bits (McMaster-Carr). The exposed brain was gently irrigated with ice-cold artificial cerebrospinal fluid and the cranial window was secured with superglue and dental cement. Mice were provided with heat, fluids, and analgesics during post-surgical recovery in accordance with institutional IACUC regulations. T-cell injections were given 15 days after tumor implantation and imaging was performed at 1 day, and 2 days after adoptive transfer of T cells.

Immunofluorescence

Mice were euthanized before being perfused transcardially with cold PBS. Brains were then removed and fixed overnight in 4% PFA–PBS before being transferred to 30% sucrose and were allowed to sink (1-2 d). Subsequently, the brains were embedded in O.C.T. Compound (Tissue-Tek; 4583; Sakura Finetek). Serial 10-µm coronal sections were then cut on freezing microtome and stored at -20 $^{0}$C. Sections were later thawed and stained overnight at 4 $^{0}$C. Primary antibodies used were: CD45 (D9M8I) XP® Rabbit mAb (Cell Signaling Technologies,

1:100), Anti–EGFRvIII, clone DH8.3 (Millipore Sigma, 1:100), Human EphA2 Alexa Fluor 700-conjugated Antibody (R&D Systems, 1:100), and Anti-IL13 receptor alpha 2 antibody (Abcam, 1:100). Secondary antibodies raised in donkey and conjugated with AlexFluor 647 were used at 4 $^0$C for two hours to detect primary labeling. Sections were stained with nuclear dye DRAQ7 (Abcam) or DAPI (Thermofisher). Images were acquired using either a Zeiss Axio Imager 2 microscope (×20 magnification) with TissueFAXS scanning software (TissueGnostics) or a Zeiss LSM 780 microscope (x20 magnification) with Zeiss Zen imaging software. Exposure times and thresholds were kept consistent across samples within imaging sessions. Single color images were used to elucidate the regions for quantification. The DAPI monochrome was used for nuclear segmentation. Colocalization of CD45 (AF647) and GFP signal with nuclear stain was done to obtain counts for GFP+ T (Strataquest software).

Assessment of Engineered T cells in vivo

For all experiments involving phenotyping of adoptively transferred engineered T cells, brain and spleen were harvested following perfusion with cold PBS. Brains were mechanically minced and treated at 37degC for 30 minutes with digestion mix consisting of Collagenase D (30mg/ml) and DNAse (10mg/ml) and Soybean trypsin Inhibitor (20mg/ml). The resulting brain homogenate was resuspended in 70% Percoll (GE Healthcare), overlaid with 30% Percoll, and then centrifuged for 30 minutes at 650g. Enriched brain infiltrating T cells were recovered at the 70-30% interface and stained with fluorescently conjugated antibodies against CD3 and CD45. Prior to staining with antibodies, cells were stained with BD Horizon Fixability Viability Stain 780 (BD Biosciences) to discriminate live from dead cells. Data was collected on Attune NxT Flow Cytometer and the analysis was performed in FlowJo software (TreeStar).

Methods References:

### SUPPLEMENTAL FIGURES:

**Fig. S1. Construction and testing of α-EGFRvIII synNotch→ α-EphA2/IL13Rα2 CAR T cells against U87 GBM, Related to Figure 1**

A Domain architectures of α-IL13Ra2/EphA2 tandem CAR, α-EphA2 CAR, and α-IL13Ra2 CAR

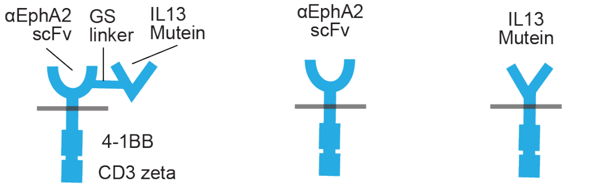


IL13 mutein: mutant form of IL13 that binds preferentially to IL13 receptor α2 (IL13Rα2)

B Relative activity of each CAR measured by in vitro killing assays of U87 GBM cell line (EphA2+ IL13Rα2+) (all constitutive expression in CD8+ T cells, NOT synNotch induced)

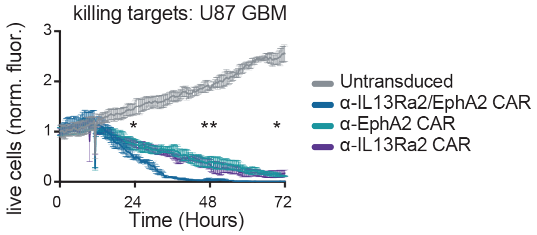


CONCLUSION: tandem CAR is more effective at in vitro killing than either single CAR

C synNotch response (expression of GFP-tagged CAR) is activated by EGFRvIII+ priming cells

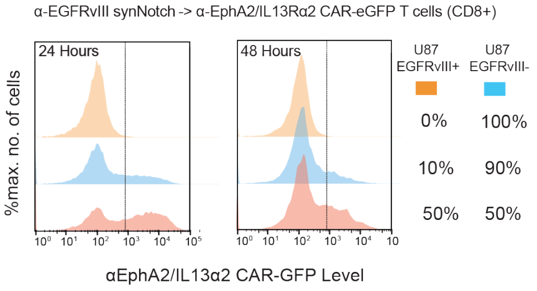


CONCLUSION: newly engineered α-EGFRvIII synNotch functions in T cells

D Gating of T cells transduced with synNotch receptor (myc-tagged) and response element (BFP tagged)

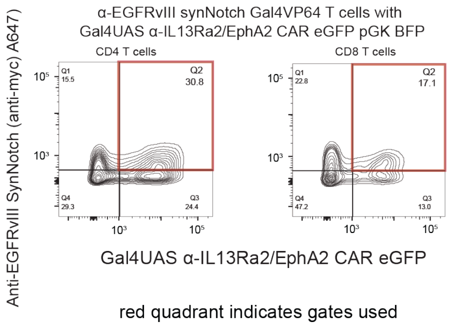


red quadrant indicates gates used

**Fig. S1. Construction and testing of α-EGFRvIII synNotch → α-IL13Rα2/EphA2 CAR T cells against U87 GBM, Related to Figure 1**

A. Domain architecture of α-IL13Rα2/EphA2 CAR, α-EphA2 CAR, and α-IL13Rα2 CAR. IL13 mutein is a mutant form of IL13 (E13K, K105R) that preferentially binds to IL13Rα2 (Krebs et al., 2014).
B. Comparing killing of U87 wild-type cells (EphA2+ IL13Rα2+) by the new α-IL13Rα2/EphA2 CAR versus T cells expressing either an α-EphA2 CAR or an α-IL13Rα2 CAR. Killing was measured using fluorescently labelled U87 cells in an IncuCyte killing assay measuring total fluorescence (live cells) over time (n=3, error bars are SEM). The tandem CAR kills more rapidly and effectively than either individual target CAR at timepoints 24, 48, and 72 hours ($p \leq .0164$; Tukey's multiple comparisons test).
C. Primary CD8+ synNotch CAR T cells described in Figure 3a were co-cultured with U87 cells described in Figure 1C. T cell priming after 24-hour and 48-hour exposure was measured by tracking induction of α-IL13Rα2/EphA2 CAR fused with a GFP reporter. FACS histograms show no induction in the absence of priming cells, and significant induction with as low as 10% priming cells (EGFRvIII-positive) (representative of at least 3 independent experiments).
D. Representative contour plots showing expression of the α-EGFRvIII synNotch Gal4VP64 receptor and the corresponding response elements regulating α-IL13Rα2/EphA2 CAR 4-1BBζ CAR GFP pGK BFP in primary CD4+ and CD8+ T cells. T cells positive for the synNotch receptor was stained through the myc-tag present on the synNotch receptor and the response element was selected based on BFP expression. The T cells in the red-boxed quadrant were sorted were sorted for in vitro and in vivo experiments.

**Fig. S2. Prime-and-kill T cells can overcome heterogeneity using a model antigen system (priming antigen: surface GFP; killing antigen CD19) in vitro.**

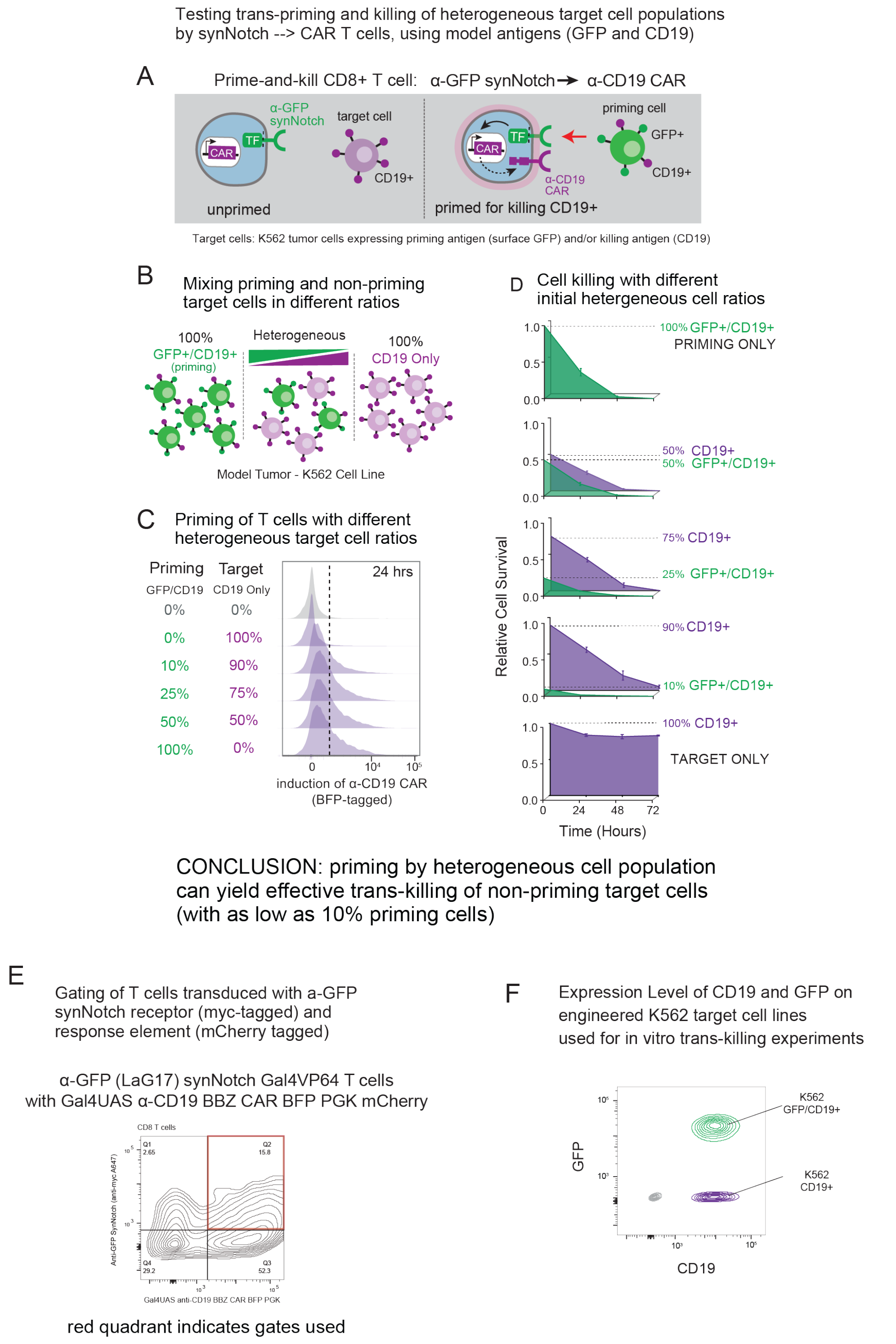

**Fig. S2. Prime-and-kill T cells can overcome heterogeneity using a model antigen system (priming antigen: surface GFP; killing antigen CD19) in vitro.**

A. Design of αGFP (Lag-17) synNotch □αCD19 4-1BBζ CAR (prime-and-kill) CD8+ T cells expressing this circuit are designed to ignore K562 target cell line expressing the killing antigen CD19 only (left), but kill cells that express both GFP and CD19 (priming and killing antigens; right).
B. We mimicked tumor heterogeneity by mixing K562 priming cells (GFP+CD19+) and target only cells (CD19+) in different priming/target cell ratios (from 0-100%).
C. Primary human CD8+ T cells expressing the circuit described in (a) were co-cultured with the K562 mixtures described in (b). T cell priming after 24-hour exposure was measured by tracking induction of CD19 CAR fused with a BFP reporter. FACS histograms show no induction in the absence of priming cells, and significant induction with as low as 10% priming cells (representative of at least 3 independent experiments).
D. Killing assays with different priming/target cell ratios. Primary CD8+ cells with the circuit described in (a) were co-cultured with K562 cells described in (b). Relative cell survival over 72 hours was quantified and showed cytotoxicity capacity of synNotch CAR T cells to overcome heterogeneous populations (n=3, error bars are SEM).
E. Representative contour plots showing expression of the αGFP synNotch Gal4VP64 receptor and the corresponding response elements regulating αCD19 4-1BBζ CAR BFP pGK mCherry in primary CD4+ and CD8+ T cells. T cells positive for the synNotch receptor was stained through the myc-tag present on the synNotch receptor and the response element was selected based on mCherry expression. The T cells in the red-boxed quadrant were sorted and used for experiments in Figure S2.
F. Flow cytometry plots showing the expression level of CD19 and GFP (Green) on dual antigen K562s and CD19 on single antigen K562s (purple) utilized for in vitro experiments.

**Fig. S3. T cells with α-EGFRvIII synNotch → α-EphA2/IL13α2 prime-and-kill circuit mediate effective and localized anti-tumor response against U87 GBM that heterogeneously express EGFRvIII in the brain. Related to Fig. 2**

Testing prime-and-kill circuit against engineered heterogeneous U87 tumors
with variable starting percentage of priming cells (EGFRvIII+)

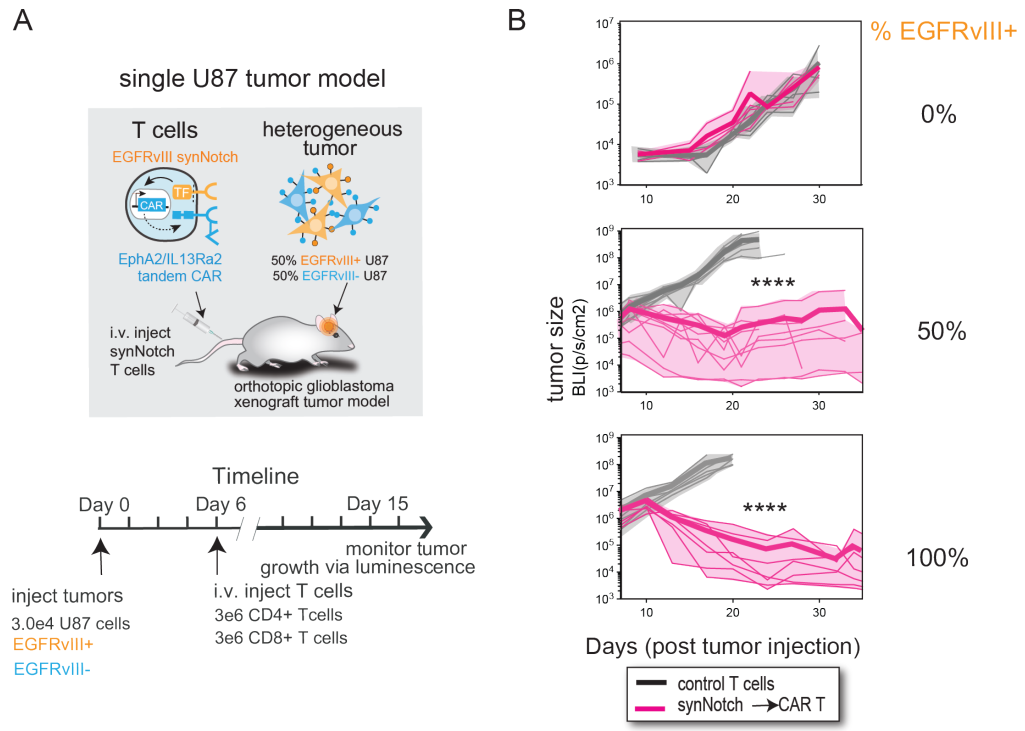


CONCLUSIONS: No in vivo tumor killing is observed when priming cells are absent.
Tumor killing in vivo is observed with 50 and 100% priming cells

C replicate of experiment shown in Fig. 2F

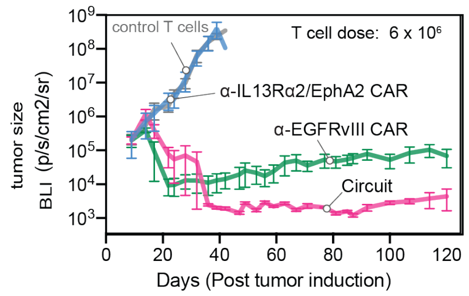


CIRCUIT: α-EGFRvIII synNotch → α-IL13Rα2/EphA2 CAR

**Fig. S3. T cells with α-EGFRvIII synNotch → α-IL13Rα2/EphA2 prime-and-kill circuit mediate effective and localized anti-tumor response against U87 GBM that heterogeneously express EGFRvIII in the brain.**

A. Immunodeficient NCG mice were orthotopically implanted in the brain with U87 GBM xenograft. Tumors contained one of the following three priming/target cell ratios: i) 100% U87 (EGFRvIII-negative) target tumor cells, ii) 50%/50% of U87-EGFRvIII-positive (priming) and EGFRvIII-negative (target) tumor cells, and iii) 100% U87-EGFRvIII-positive (priming) cells. Tumor cells were engineered to express luciferase to allow for tracking of tumor size. Six days following tumor implantation, the mice were infused intravenously with 3 million each of CD4+ and CD8+T cells. T cells expressed either: i) no construct (non-transduced control) or ii) α-EGFRvIII synNotch→ α-IL13Rα2/EphA2 CAR circuit.
B. Tumor size was determined by longitudinal bioluminescence imaging. Individual traces for each animal are shown with thin lines, while average is shown with the thick line. Negative control treatment with non-transduced T cells is shown in black, prime-and-kill CAR T cell treatment is shown in pink. Prime-and-kill CAR T cells have no impact on tumors that lack EGFRvIII priming (left panels, n=5), but show significant improvement in reducing tumor size (**** $p< 0.0001$, t test).
C. Replicate of the data shown in Figure 2D, F. NCG mice were orthotopically implanted in the brain with GBM6 PDX cells. Tumor cells were engineered to express mCherry and luciferase to allow for tracking of tumor size. Ten days following tumor implantation, the mice were infused intravenously with 3 million each of CD4+ and CD8+T cells. T cells expressed either: i) no construct ( control) (n=5), ii) α-EGFRvIII synNotch→ α-IL13Rα2/EphA2 CAR circuit (n=6), iii) constitutively expressed α-EGFRvIII CAR (n=5), or iv) constitutively expressed α-IL13Rα2/EphA2 tandem CAR (n=5). Time course tracking of tumor size, determined by longitudinal bioluminescence imaging. Negative control treatment with non-transduced T cells is shown in black (n=5), prime-and-kill CAR circuit treatment is shown in pink (n=6), conventional α-EGFRvIII CAR treatment is shown in green line (n=6), and constitutive α-IL13Rα2/EphA2 tandem CAR in blue (n=5). Mice treated with constitutive α-IL13Rα2/EphA2 tandem CAR failed to control tumor growth. Treatment with conventional α-EGFRvIII CAR T cells resulted in early tumor regression followed by recurrence in all mice (n=6). In contrast, all mice treated with the prime-and-kill CAR T cells showed complete clearance of tumor ($p< 0.0001$ t-test untransduced vs prime-and-kill CAR T cells). Error bars represent mean ± SEM of 5-6 individual mice from one experiment

**Fig. S4. Testing α-EGFRvIII synNotch → α-IL13Rα2/EphA2 CAR T cells against GBM6, Related to Figure 3**

A

GBM6 cells activate α-EGFRvIII synNotch

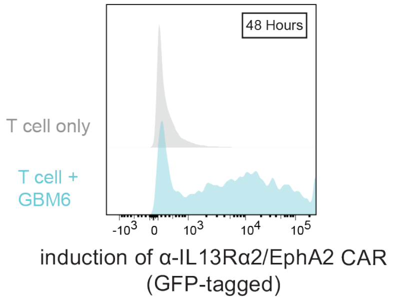


C

GBM6 PDX cells show intrinsic heterogeneity of EGFRvIII expression

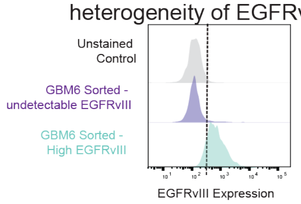


CONCLUSIONS: we can sort GBM6 cells to separate high EGFRvIII expression cells from cells with undetectable EGFRvIII expression

B

GBM6 cells are killed effectively in vitro by T cells with α-EGFRvIII synNotch-> α-IL13Rα2/EphA2 CAR circuit

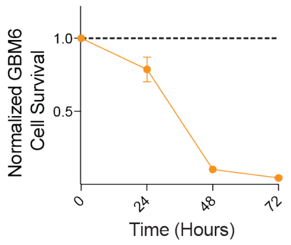


CONCLUSIONS: EGFRvIII synNotch to CAR circuit is primed by GBM6 PDX cells, and yields complete killing of GBM6 cells in vitro

D

Killing of GBM6 cells by EGFRvIII primed T cells is dependent on EGFRvIII expression levels

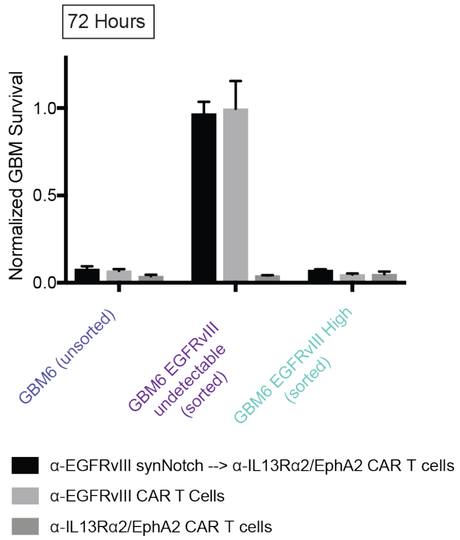


CONCLUSIONS: by sorting high and low EGFRvIII expressing GBM6 cells, we find that EGFRvIII expression is necessary for killing by EGFRvIII primed T cells

**Fig. S4. Testing α-EGFRvIII synNotch → α-IL13Rα2/EphA2 CAR T cells against GBM6, Related to Figure 3**

A. Primary CD8+ α-EGFRvIII synNotch → α-IL13Rα2/EphA2 CAR T cells were co-cultured with or without GBM6 cells at 1:1 ET ratio. T cell priming after 48-hour exposure was measured by tracking induction of α-IL13Rα2/EphA2 CAR fused with a GFP reporter. FACS histograms show no induction in the absence of GBM6 cells, and significant induction with GBM6 cells (representative of at least 3 independent experiments).
B. Killing assays with primary CD8+ α-EGFRvIII synNotch → α-IL13Rα2/EphA2 CAR T cells with the circuit were co-cultured with GBM6 cells at 1:1 ET ratio. Relative cell survival over 72 hours was quantified and showed cytotoxicity capacity of synNotch CAR T cells to overcome GBM6 cell populations (n=3, error bars are SEM).
C. GBM6 cell were sorted for varying level of EGFRvIII expression and evaluated by flow cytometry post-sort. Gray represents unstained control.
D. Killing assays with primary CD8+ α-EGFRvIII synNotch →α-IL13Rα2/EphA2 CAR T cells, α-EGFRvIII CAR T cells, or α-IL13Rα2/EphA2 CAR T cells co-cultured with either none, high, or unsorted expression of EGFRvIII. Relative cell survival over 72 hours was quantified and showed inability of α-EGFRvIII synNotch → α-IL13Rα2/EphA2 CAR T cells and α-EGFRvIII CAR T cells to kill GBM6 cell population that did not express any EGFRvIII antigen (n=3, error bars are SEM).

**Fig. S5. Testing α-EGFRvIII synNotch→CAR circuit T cells display reduced differentiation state comparedto standard constitutive CARs, Related to Figure 3**

A

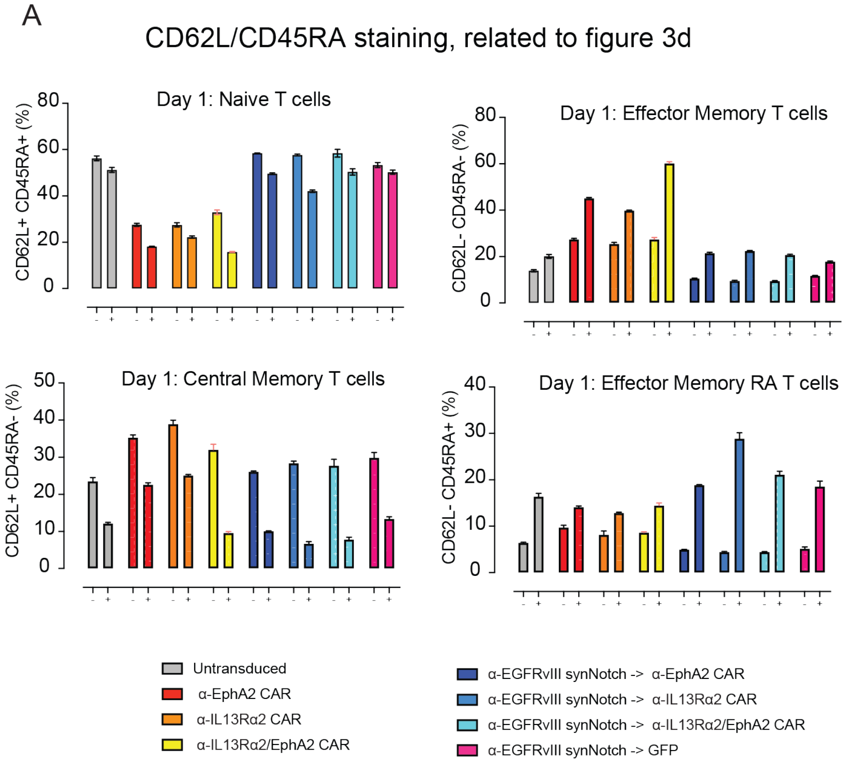


B

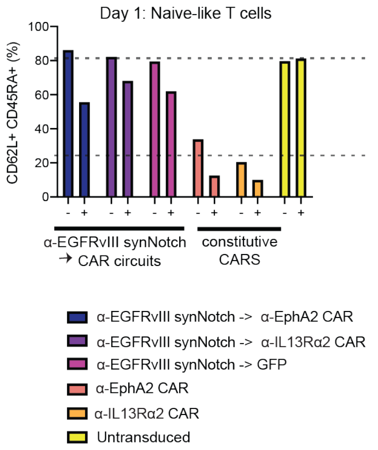


Biological replicate of cell differentiation of different circuit examined in Fig. 3d

comparing state of constitutively expressed single CARs (EphA2 CAR or IL13Ra2 CAR) with synNotch induced versions

CONCLUSION: synNotch induction of single CAR constructs (which are in clinical trials) significantly increases percentage of naive-like cells

**Fig. S5. Testing α-EGFRvIII synNotch → CAR circuit T cells display reduced differentiation state compared to standard constitutive CARs, Related to Figure 3**

A. Surface expression of CD45RA and CD62L to distinguish T memory stem cells (CD45RA+CD62L+), central memory cells (CD45RA−CD62L+), effector memory cells (CD45RA−CD62L−), and effector memory RA cells (CD45RA+CD62L-) in all relevant CAR T cells for this study, including constitutive α-EphA2 CAR, constitutive α-IL13Rα2 CAR, and constitutive α-IL13Rα2/EphA2 tandem CAR, as well as the α-EGFRvIII synNotch primed versions of these circuits (n=3). (Same experiment as 3d)
B. Percentage of CD62L+CD45RA+ phenotype cells in all relevant CAR T cells for this study, including constitutive α-EphA2 CAR, constitutive α-IL13Rα2 CAR, and constitutive α-IL13Rα2/EphA2 tandem CAR, as well as the α-EGFRvIII synNotch primed versions of these circuits. Data is shown with and without GBM6 cells (after 24 hours). Bar graph show the percent of T cell population with memory stem-cell phenotype (Biological Replicate of 3d).

**Fig. S6. Local brain-specific prime-and-kill CAR T cells mediate effective anti-GBM responses**

A FACS analysis of α-MOG synNotch --> α-IL13Ra2/EphA2 CAR T cells (Fig. 4.A-F recovered from spleen or brain (day 6 after T cell injection)

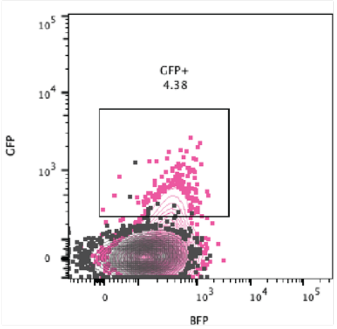


MOG circuit (BFP tagged)
priming: expression of GFP-tagged CAR

spleen, MOG circuit (BFP+)
brain, MOG circuit (BFP+)

CONCLUSION: MOG synNotch circuit T cells show priming in brain but not in spleen

B Analysis of additional mice: local priming of α-MOG synNotch --> α-IL13Ra2/EphA2 CAR T cells (related to Fig. 4D)

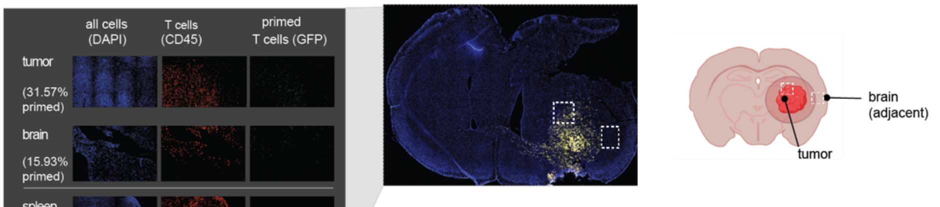


GBM6 tumor (6 days after T cell injection)

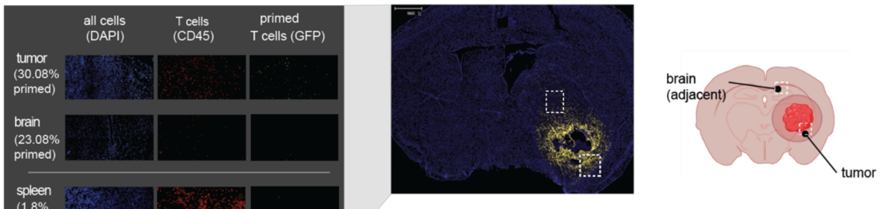


GBM6 tumor (6 days after T cell injection)

**Fig. S6. Local brain-specific prime-and-kill CAR T cells mediate effective anti-GBM responses**

A. Flow cytometry of MOG synNotch CAR T cells isolated from brain and spleen of GBM6-bearing mouse demonstrates presence of GFP+ primed T cells in the brain. SynNotch CAR T cells express BFP constitutively. T cells were pregated on CD3 and CD45.
B. Replicates of data shown in Figure 4D. Observation of T cell priming in the brain and tumor. GBM6 tumor-bearing mice were euthanized six days after MOG SynNotch →CAR circuit T cell infusion. Representative confocal fluorescent microscopy of sections obtained from circuit T cell-treated mice reveals primed T cells (GFP+ indicates priming; red indicates human T cells identified with hCD45 stain) in the tumor (yellow) and in the infiltrative parts of the tumor. Quantification of GFP+ T cells in tumor using nuclear segmentation reveals greater number of GFP+ T cells in tumor core than in periphery. Circuit CAR T cells (red) in the spleen do not express GFP. Insets (single stained images, scale bar: 50µm) are enlargements of outlined regions in the main image.

#### Fig. S7. Construction and testing of α-CDH10 synNotch ➔ α-IL13Rα2/EphA2 CAR T cells

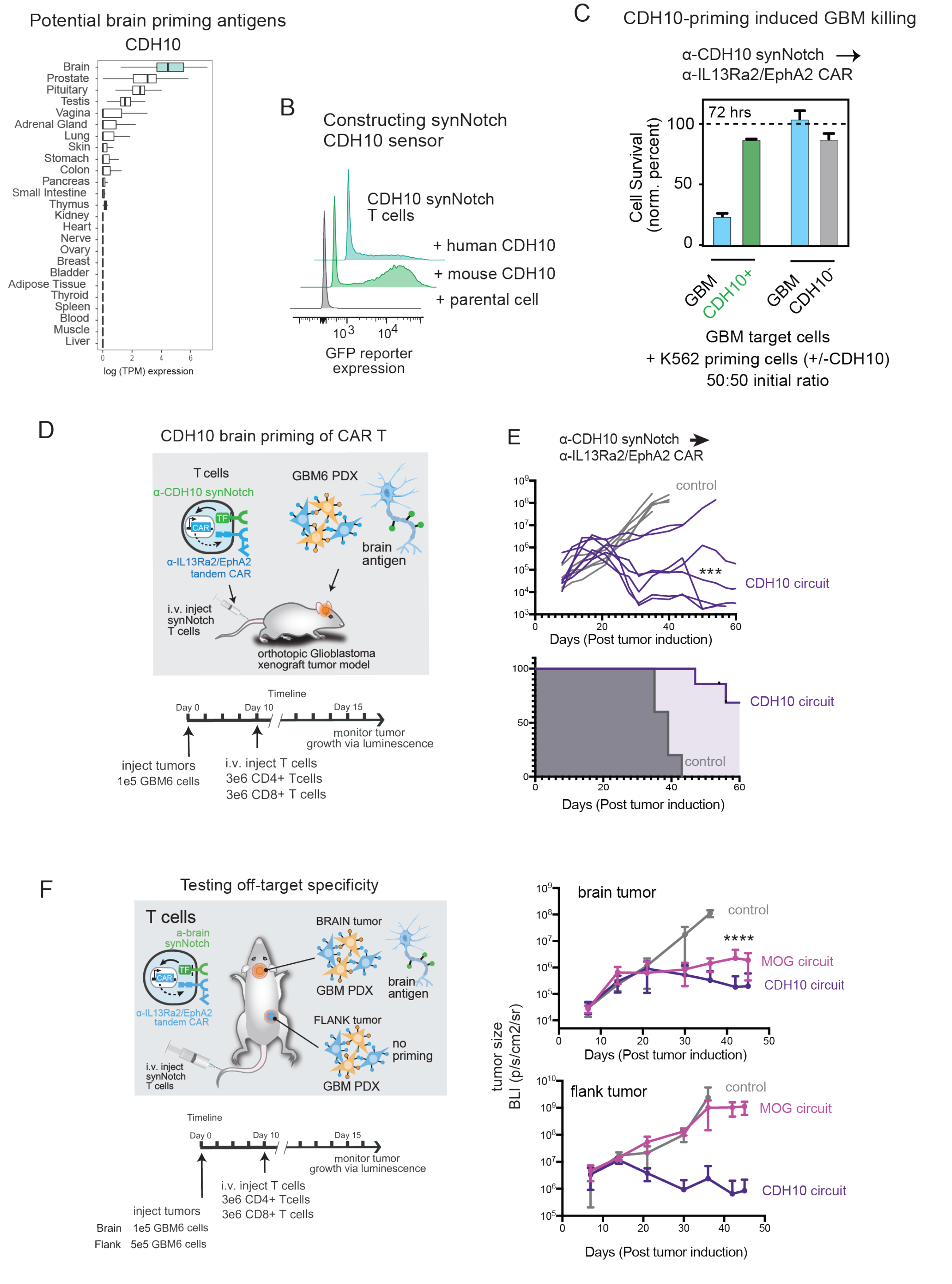


CONCLUSIONS: MOG priming does not induce killing of flank GBM6 tumor but CDH10 priming does. Thus MOG appears to be more precise brain priming antigen than CDH10.

**Fig. S7. Construction and testing of α-CDH10 synNotch → α-IL13Rα2/EphA2 CAR T cells**

A. Box and whisker plots showing tissue specific expression of CDH10 across a subset of tissue samples in GTEx v7. Units shown are log scaled normalized RNAseq counts (Transcripts Per Million) taken from GTEx portal v7 (https://gtexportal.org/).
B. Primary CD8+ α-CDH10 → GFP PGK BFP T cells were co-cultured with either parental K562 or K562 transduced to express mouse CDH10 or human CDH10. T cell priming after 48-hour exposure was measured by induction of GFP reporter. FACS histograms show induction of GFP reporter only in the presence of mouse CDH10+ or human CDH10+ K562 and not K562 parental cells (representative of 3 experiments).
C. Primary CD8+ α-CDH10 synNotch → α-IL13Rα2/EphA2 CAR T cells were co-cultured with U87 cells and K562 expressing or not expressing mouse CDH10. Relative cell survival over 72 hours was quantified and showed cytotoxicity capacity of α-CDH10 synNotch → α-IL13Rα2/EphA2 CAR T cells to kill U87 cells only when priming cells are expressing mouse CDH10 (n=3, error bars are SD). Significant cytotoxicity in the U87 co-cultured with K562 mouse CDH10 was observed (*** $p < 0.001$; t test) compared to the U87 co-cultured with K562 Parental. Cell population ratio: 1:1:1, 1e4 cell each.
D. Immunodeficient NCG mice were orthotopically implanted in the brain with GBM6 patient-derived xenograft cells. Tumor cells were engineered to express mCherry and luciferase to allow for tracking of tumor size. Ten days following tumor implantation, the mice were infused intravenously with 3 million each of CD4+ and CD8+T cells. T cells expressed either: i) no construct (non-transduced control) or ii) α-CDH10 synNotch→ α-IL13α2/EphA2 CAR circuit (n=7).
E. Tumor size (top row) and survival (bottom row) over time. Tumor size was determined by longitudinal bioluminescence imaging. Negative control treatment with non-transduced T cells is shown in grey, prime-and-kill CAR circuit treatment is shown in purple. Compared to non-transduced treatment group, mice treated with the α-CDH10- prime-and-kill CAR T cells (5 out of 7 mice) showed sustained anti-tumor response ($p<0.001$ by t test with Holm-Sidak correction for multiple comparisons) ($p <0.0001$ Log-rank (Mantel-Cox) test.
F. Evaluating tissue specificity of CDH10 and MOG priming circuits. GBM6 PDX tumor cells were implanted.in the brain and flank of NCG mice. Both tumors express the killing antigens (EphA2 and IL13Rα2) however the expression of priming antigen MOG/CDH10 is ideally restricted to the brain. Mice were treated one-time (10 days after tumor implantation) with intravenous infusion of non-transduced (n=5), or α-MOG-prime-and-kill CAR T cells (n=6), or (iii) α-CDH10-prime-and-kill CAR T cells (n=5). Tumor size was measured by luciferase luminescence. α-MOG-prime-and-kill CAR T cells are shown in pink, α-CDH10-prime-and-kill CAR T cells are shown in purple while the non-transduced control T cells are shown in gray. Significant suppression in the size of brain tumor was observed in prime-and-kill CAR T cell treated mice (**** $p < 0.0001$; t test) while the flank tumor grew at the same rate as in the mice treated with non-transduced T cells. Importantly MOG priming circuit does not induce killing of flank tumor, but the CDH10 priming circuit does, indicating that MOG is a more specific brain priming antigen.

**Supplemental Movies**

**Movie S1.**

Time-lapse analysis of 100%, 90% or 50% U87 EGFRvIII-negative with corresponding U87 EGFRvIII-positive cells 0%, 10% or 50% (total of 10K U87 cells) were co-cultured with primary CD8+ human T cells with the α-EGFRvIII synNotch → α-IL13Rα2/EphA2 CAR (50K T cells). EGFRvIII-positive priming cells are colored in yellow, while EGFRvIII-negative target cells are colored in blue (T cells are unlabeled). Fluorescent images were obtained every 15 minutes over 3 days using an IncuCyte Live-Cell Imaging System (Essen Bioscience, Ann Arbor, MI).

**Movie S2.**

Intravital imaging (Two-photon in vivo laser scanning) of circuit CAR T cells show dynamic priming within the GBM6 xenograft tumor. Tumors were implanted at a depth of 3mm below the right frontal cortex and cranial windows were implanted; tumor cells, orange; unprimed circuit CAR T cell, blue; primed circuit CAR T cell, green.
